# A New Approach to Symmetric Registration of Longitudinal Structural MRI of the Human Brain

**DOI:** 10.1101/306811

**Authors:** Babak A. Ardekani, for the Alzheimer’s Disease Neuroimaging Initiative

**Author notes:** Center for Brain Imaging and Neuromodulation, The Nathan S. Kline Institute for Psychiatric Research, 140 Old Orangeburg Road, Orangeburg, NY 10962, USA. Data used in preparation of this article were obtained from the Alzheimer’s Disease Neuroimaging Initiative (ADNI) database (adni.loni.usc.edu). As such, the investigators within the ADNI contributed to the design and implementation of ADNI and/or provided data but did not participate in analysis or writing of this report. A complete listing of ADNI investigators can be found at: http://adni.loni.usc.edu/wp-content/uploads/how_to_apply/ADNI_Acknowledgement_List.pdf.

## Abstract

This technical report presents the Automatic Temporal Registration Algorithm (ATRA) for symmetric rigid-body affine registration of longitudinal *T*_1_-weighted three-dimensional magnetic resonance imaging (MRI) scans of the human brain. This is a fundamental processing step in computational neuroimaging. The notion of leave-one-out consistent (LOOC) landmarks with respect to a supervised landmark detection algorithm is introduced. An automatic algorithm is presented for identification of LOOC landmarks on MRI scans. Using this technique, multiple sets of LOOC landmarks (around 150) are identified on each of the volumes being registered. Then, a Generalized Orthogonal Procrustes Analysis of the identified landmarks is used to find a rigid-body transformation of each volume into a common space where the transformed volumes are precisely aligned. In addition, a new approach is introduced for quantitative assessment of registration accuracy in the absence of a gold standard. Qualitative and quantitative evaluations of ATRA registration accuracy are performed using 2012 volumes from 503 subjects (4 longitudinal volumes/subject) from the Alzheimer’s Disease Neuroimaging Initiative database, and on a further 120 volumes acquired from 3 normal subjects (40 longitudinal volumes/subject). The algorithm is symmetric, in the sense that any permutation of the input volumes does not change the resulting transformation matrices, and unbiased, in the sense that all volumes undergo one and only one interpolation operation, which precisely aligns them in a common space. There is no interpolation bias and no reference volume. All volumes are treated exactly the same. The algorithm is fast and highly accurate. The software is publicly available.

## 1 Introduction

Medical image registration is the process of estimating a one-to-one mapping between physically corresponding points within the fields-of-view (FOVs) of a pair of scans. The scans can be from different imaging modalities (e.g., PET and MRI) obtained from the same individual (Ardekani et al., 1995), from the same modality (e.g., MRI-MRI) obtained from different individuals (Klein et al., 2009), or from the same modality and individual (e.g., longitudinal MRI) (Hajnal et al., 1995). Most registration algorithms between pairs of images have been designed asymmetrically (Nestares and Heeger, 2000; Holland and Dale, 2011) whereby, somewhat arbitrarily, one scan is taken as the *target* (also referred to as the *reference* or *template* scan) and the other as the *source* (also referred to as the *moving* or *subject* scan). The objective is to find a transformation from within a class of transformations that, when applied to the source image, would match it as closely as possible to the target image in some sense. A problem with this approach is that the resulting registrations are not necessarily *inverse-consistent*, that is, if the roles of the target and source images were reversed, the algorithm would not necessarily yield the *inverse* transformation. Another major problem with asymmetric registration is the so-called interpolation bias, whereby the target image remains fixed while the source image undergoes an interpolation with associated smoothing. This can be detrimental, for example, in applications where a pair of serial MRI volumes are matched in order to detect subtle atrophic changes over time (Liu et al., 2003).

To resolve these issues, researchers have been working on designing inverse-consistent and unbiased registration techniques (Smith et al., 2002; Johnson and Christensen, 2002; Reuter et al., 2010; Ardekani et al., 2016) between pairs of scans. There have also been efforts to extend symmetric registration to the problem of matching multiple (greater than two) scans (Reuter et al., 2012; Ashburner and Ridgway, 2012). Although, to avoid high computational cost, Reuter et al. (2012) suggest a compromised implementation, whereby first the registration of each image to a randomly selected image is computed. Therefore, this approach is not truly symmetric; it merely randomizes the asymmetry. The method described by Ashburner and Ridgway (2012) is similarly computationally expensive.

In this technical report, we introduce the Automatic Temporal Registration Algorithm (ATRA), a new approach to rigid-body affine registration which is truly symmetric regardless of the number of scans being registered and computationally efficient. Although ATRA has been *trained* for the registration of *T*_1_-weighted three-dimensional (3D) structural MRI brain scans (volumes), the same process of training can in principle be applied to registering other modalities (e.g., *T*_2_-weighted volumes). Rigid-body registration of serial volumes acquired from the same individual at different times is a fundamental processing step in computational neuroimaging. Multiple independent scans may be acquired during the same scanning session or longitudinally separated in time by days, months or years. For example, several structural scans with short acquisition times are sometimes obtained in the same scanning session to reduce motion artifacts (Marcus et al., 2007). The volumes are then retrospectively registered and averaged to obtain a single volume with higher signal-to-noise ratio (SNR). The structural MRI protocol in the Alzheimer’s Disease Neuroimaging Initiative (ADNI) calls for back-to-back structural MRI scans within the same session to reduce the number of image acquisition sessions that must be repeated due to poor image quality (Jack et al., 2008). Registration of the back-to-back scans may be necessary in some applications, e.g., in studying the reproducibility of automated hippocampal volumetry software (Bartel et al., 2017).

Comparison of longitudinally acquired scans is a powerful tool for diagnosis, monitoring disease progression, evaluation of treatments and research. For example, registration and subtraction of serially acquired brain MRI pre- and post-surgery significantly increases the sensitivity of radiologists in finding new ischemic lesions following cardiac surgery (Patel et al., 2017). The subtraction technique is also extremely useful in monitoring new lesion activity in multiple sclerosis (Sweeney et al., 2013). Registration of longitudinal MRI from the same individual is also a prerequisite for subsequently more sophisticated methods that are designed to automatically measure regional or global brain volume change (Freeborough and Fox, 1997; Smith et al., 2002).

A problem with using the rigid-body model for the registration of longitudinal data is that there are regions in the imaging FOV where this model does not hold. Firstly, there are moving parts such as the jaw, eyes, tongue, neck and scalp that cannot be expected to be rigid. Secondly, there may be subtle changes in the structure of the brain over time, such as ventricular enlargement, expanding or new white matter lesions, and brain atrophy. Therefore, registration methods have to be designed to handle departures from the rigid-body motion model. For this purpose, Reuter et al. (2012) proposed using robust estimation methods designed to automatically reduce the influence of outlier regions from the registration process. Ashburner and Ridgway (2012) propose a scheme which combines non-linear diffeomorphic and rigid-body registrations along with correction for the intensity inhomogeneity artifacts.

Here we postulate that there exist “stationary landmarks” within the brain which we refer to as *anchor points* whose relative positions remain constant over the time periods typically covered in longitudinal imaging. Well known examples of these points are the intersections of the anterior and posterior commissures with the mid-sagittal plane of the brain. We present a method for automatically and rapidly identifying a large number of these points (about 150) across all brain volumes being registered and finding rigid-body transformations that would match these points on a common space in a fully symmetric manner. We will show this method to be highly accurate.

In the material presented in this paper, we make a distinction between landmark *identification* and landmark *detection*. The distinction is best described in the context of a *supervised* landmark detection algorithm. In the *training* phase, the algorithm is provided with a set of prototype landmarks on multiple training volumes. In most previously published work, these prototype landmarks are *identified* manually on the training set of volumes by expert raters (Ardekani and Bachman, 2009; Ghayoor et al., 2017). This is a time-consuming, tedious, and error-prone process for which we use the term *landmark identification*. On the other hand, once the algorithm is trained, we refer to the process of using the algorithm for estimating the location of the same landmark on a test volume as *landmark detection*.

A problem with manual landmark identification is that it is very difficult for humans to define large numbers of homologous points that can be called landmarks. Usually it is difficult to define more than a few dozen different landmarks across the brain. Even when a landmark is clearly defined, it is often an extremely difficult task to locate it manually across multiple individuals. Furthermore, a point that humans can recognize and reliably locate as a brain landmark is not necessary ideal for automatic detection.

A major contribution of this work is an algorithm for automatic landmark identification which can be used for training landmark detection algorithms. Furthermore, since the algorithm is fast and fully automatic, it can be used to identify a large number of homologous points across scans that can then be used for image registration.

Specific contributions of this paper are as follows: (1) the notion of leave-one-out consistent (LOOC) landmarks is introduced; (2) a simple but very useful algorithm for automatically and rapidly identifying multiple sets of LOOC landmarks is presented; (3) a truly symmetric and computationally efficient intra-subject intra-modality image registration algorithm based on LOOC landmarks is presented; (4) a novel method for assessing the accuracy of image registration in the absence of a gold standard is introduced; and finally (5) an implementation of the algorithm (ATRA) has been made freely available to the research community (www.nitric.org/projects/art).

## 2 Methods

Assume that we have a set of 3D structural MRI volumes *V*^(*m*)^ (*m* = 1,2 …, *M*) acquired from the same individual at different times. It is useful to think of these volumes as functions *V*^(*m*)^: ℝ^3^ × {1} → ℝ that map real-world coordinates [*x y z* 1]^*T*^ in the imaging FOV, expressed in homogeneous coordinates notation as four dimensional column vectors, to real numbers representing the image values. The objective of ATRA is to find *M* corresponding rigid-body transformations *T*^(*m*)^: ℝ^3^ × {1} → ℝ^3^ × {1} that transform the *M* volumes to a common space in which they are precisely registered. ATRA is symmetric in the sense that any permutation of {*V*^(1)^, *V*^(2)^,…, *V*^(*M*)^} results in the same permutation of {*T*^(1)^,*T*^(2)^,…,*T*^(*M*)^} (Ashburner and Ridgway, 2012). In the ensuing discussion, we will drop the superscript *m* whenever convenient.

The affine transformations *T* are expressed as 4 × 4 matrices of the form:

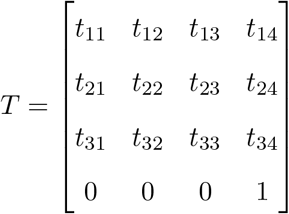

along with the rigid-body constraint |*T*| = ±1. These transformations map real-world coordinates *q* = [*x y z* 1]^*T*^ in the domain (FOV) of *V* to real-world positions *T*(*q*) = *w* in the common space. Accordingly, we can denote a transformed volume in the common space by the composite function 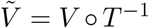 defined by 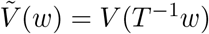. In practice, since the MRI FOV is discretized, evaluations of function *V* at arbitrary points *T*^−1^*w* requires spatial interpolation. Throughout this work, we have used linear interpolation for this purpose.

In ATRA, the solution transformations are expressed as: 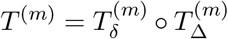, where the transformation 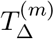 is found separately for each volume using intrinsic characteristics of the volume and represents a coarse rigid-body transformation to an intermediate standard space. The algorithm for obtaining 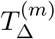 is described in Section 2.1. The transformations 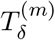 are found simultaneously and symmetrically for all *M* volumes to correct for small residual misalignments that remain between volumes following transformation to the intermediate standard space. The algorithm for obtaining 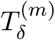 is presented in Section 2.7.

### 2.1 Alignment to standard orientation

In this section we present a fast and fully automated algorithm for computing *Τ*_Δ_ which transforms an arbitrarily oriented volume *V* to an intermediate standard orientation. The transformation is computed independently for each of the *M* volumes to be registered. The algorithm for computing *Τ*_Δ_ is based on intrinsic characteristics of the MRI volume *V* alone and does not require a reference volume. Without loss of generality, we defined the standard space such that:

1. The *z* = 0 plane closely coincides with the the brain’s mid-sagittal plane (MSP).
2. The origin (FOV center) is approximately the mid-point between the intersection points of the anterior commissure (AC) and the posterior commissure (PC) on the MSP.
3. The *x* axis is on the MSP, approximately parallel to the AC-PC line and points posteriorly.
4. The *y* axis is on the MSP, perpendicular to the *x* axis and points inferiorly.
5. The *z* axis is perpendicular to the MSP and points from subject’s right to left.

We refer to this as the posterior-inferior-left (PIL) orientation. The *z* = 0 plane of an MRI volume after transformation to PIL space is shown in Fig. 1.

**Figure 1:**
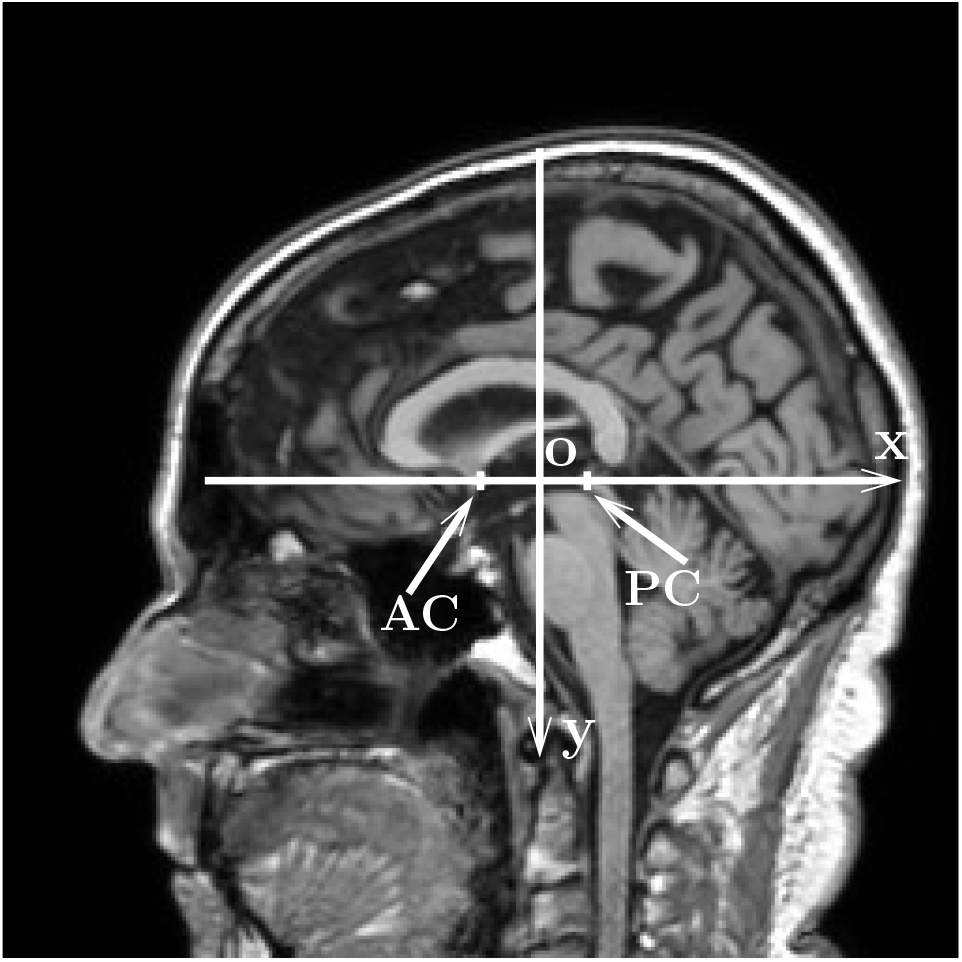
The *z* = 0 plane of a volume after transformation to PIL space. In this standard space, the *x* axis points to the posterior direction, the *y* axis points to the inferior direction, and the *z* axis (not shown) points from subject’s right to left.

Transformation of an arbitrarily oriented volume to the PIL standard space is accomplished in three steps as shown in Fig. 2. Each step involves a rigid-body transformation. The final transformation is obtained by combining the three transformations. In step 1, the mid-sagittal plane (MSP) is estimated automatically using the algorithm described by Ardekani et al. (1997). Using the estimated MSP, a transformation matrix *T_msp_* is determined so that the *z* = 0 plane of the transformed volume 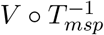 coincides with the estimated MSP.

**Figure 2:**
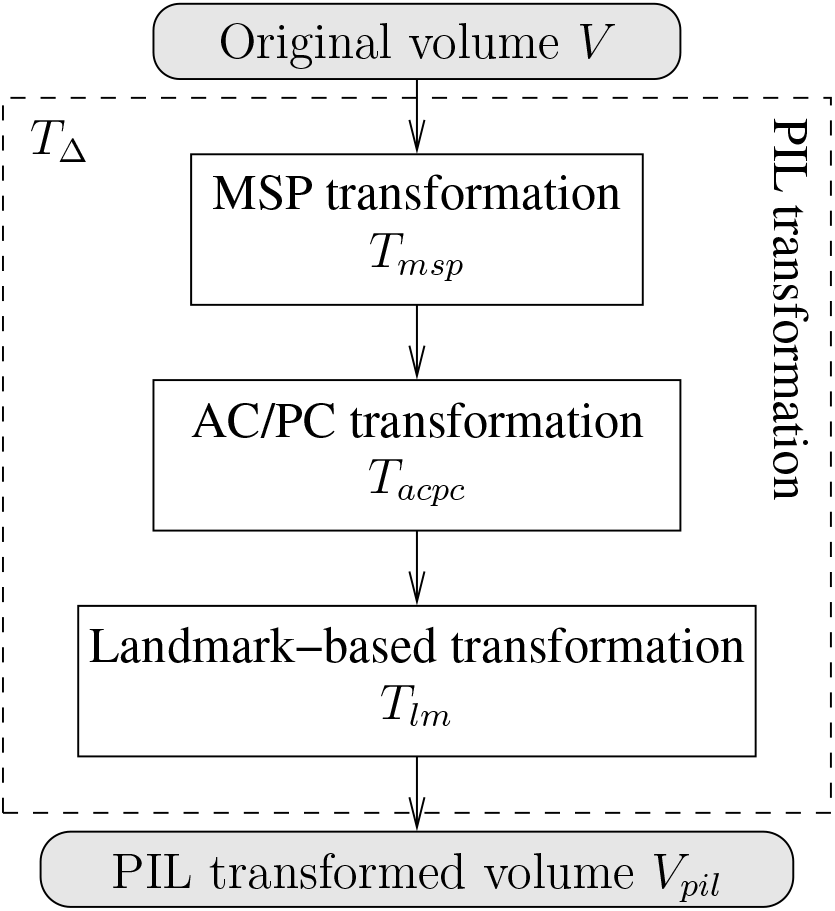
Flowchart for rigid-body transformation of an arbitrarily orientated volume *V* to the standard PIL space. The transformation is comprised of three steps that are combined into a single transformation *Τ*_Δ_ = *T*_*lm*_ ○ *T_acpc_* ○ *T_msp_* to obtain the transformed volume 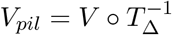.

In step 2, the mid-sagittal intersection points of the AC and the PC are automatically detected on the MSP using the algorithm described by Ardekani and Bachman (2009). Based on this information, a second rigid-body transformation is obtained which maps the center of the transformed volume’s FOV to the mid-point between the detected AC and PC points, and rotates the transformed volume so that the *x* axis points posteriorly parallel to the AC-to-PC line and the *y* axis points inferiorly perpendicular to the AC-to-PC line. The corresponding transformation matrix is denoted by *T_acpc_* (Fig. 2). Following this step, a transformed volume *V* ○ (*T_acpc_* ○ *T*_*msp*_)^−1^ would be approximately in the desired PIL orientation. However, since the AC and PC points are only approximately an inch apart on the MSP, and because the accuracy of automated AC/PC detection algorithms has been reported to be on the order of 1 mm (Ardekani and Bachman, 2009; Bhanu Prakash et al., 2006; Verard et al., 1997; Liu and Dawant, 2015), errors in AC/PC detection could result in a large variance in the head’s pitch angle (Arndt et al., 1996; Evans et al., 1992; Li et al., 2003).

Therefore, in step 3, to stabilize the PIL transformation further, eight additional landmarks {*q*_1_, *q*_2_,…, *q*_8_} are located automatically on the MSP and a third rigid-body transformation *T_lm_* is computed so that the 8 landmarks are mapped as closely as possible to 8 corresponding previously determined target locations 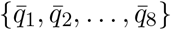 (Arun et al., 1987). More precisely, *T_lm_* is found as follows:

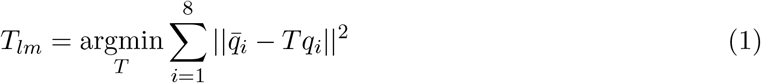

subject to the constraint |*T*_*lm*_| = 1. Methods used for identification and detection of landmarks *q_i_* and their target locations 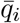 are described in Sections 2.4 and 2.5.

To summarize, given an arbitrarily oriented volume *V*, ATRA fully automatically and rapidly determines a rigid-body transformation given by *Τ*_Δ_ = *T_lm_* ○ *T_acpc_* ○ *T_msp_* such that 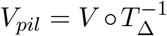 is in the standard PIL orientation. For short, we will refer to the procedure described in this section as the *PIL transformation*. As an example, Fig. 3 shows the *z* = 0 plane after PIL transformation of 4 longitudinal volumes along with the projections of the eight landmarks {*q*_1_, *q*_2_,…, *q*_8_} detected independently for each volume.

**Figure 3:**
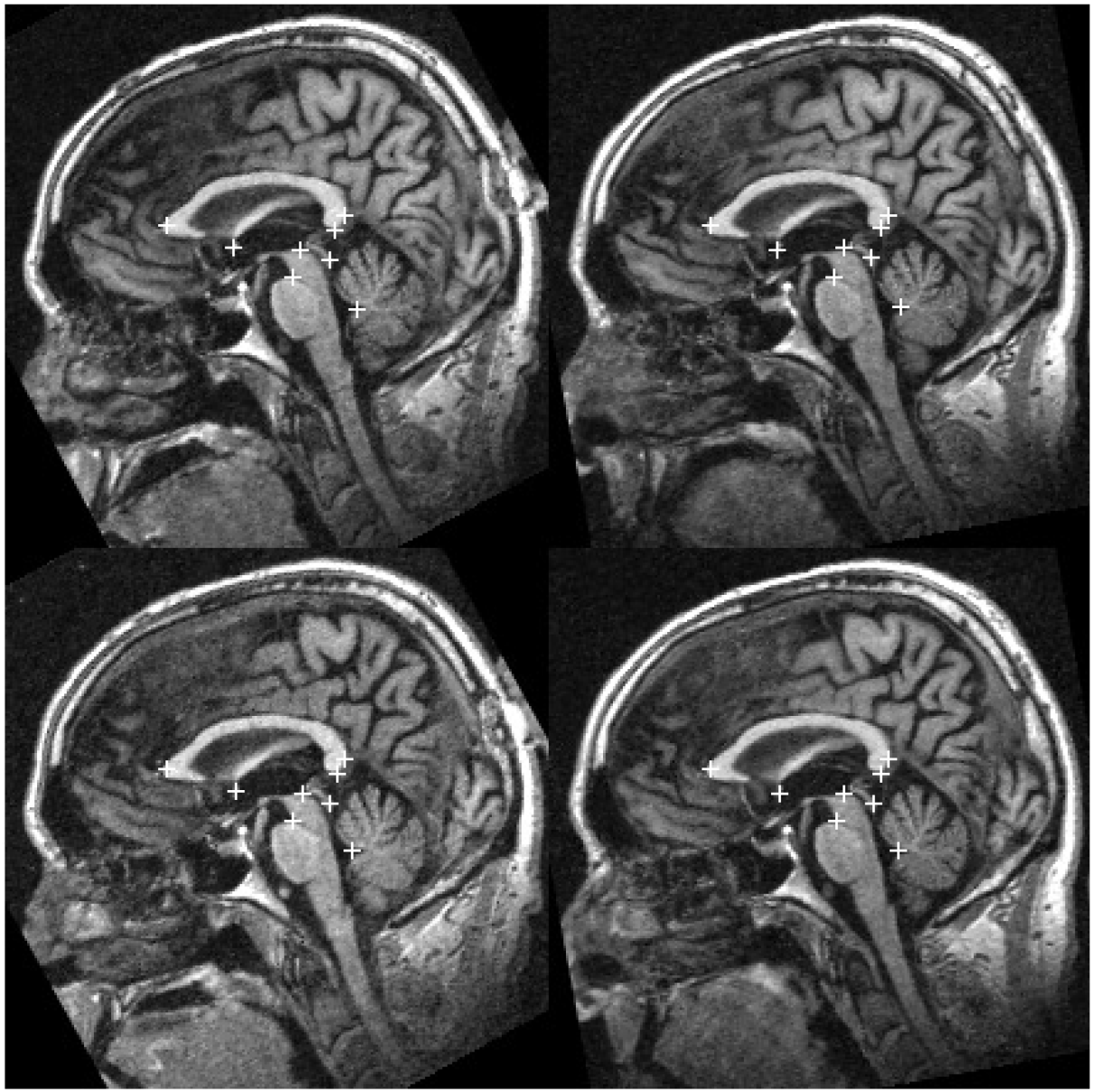
The *z* = 0 plane in four longitudinal volumes acquired from the same individual after PIL transformation 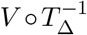. In each volume, the “+” marks indicate projections of the eight landmarks {*q*_1_, *q*_2_,…, *q*_8_} detected independently for each volume. Note that after PIL transformation of the 4 volumes a coarse registration has been achieved. However, small residual misalignments remain that will be corrected by applying *T*_*δ*_.

### 2.2 Intra-cranial space probability map

The non-brain regions in MRI volumes are problematic for brain image registration algorithms. For example, there are moving parts such as the jaw, eyes and neck that cannot be modeled by rigid-body motion. Subcutaneous fat has much shorter *T*_1_ relaxation time than brain tissue which would result in high signal intensities that could dominate computation of similarity measures such as sum of squared differences between voxel intensities. For these and other reasons, such as computation efficiency, it is very beneficial to remove or reduce the influence of the non-brain regions during image processing. To address this problem, Reuter et al. (2010) used robust statistics to automatically reduce the influence of outlier voxels which mostly reside in non-brain regions.

In this work, we employ a very simple and effective alternative method for limiting the image processing to the intra-cranial space. In an offline processing step, we utilized *M* volumes acquired from *different* individuals denoted by *V*^(*m*)^ (*m* = 1,2,…,*M*). In each volume, we isolated the intra-cranial region using the *brainwash* module of the Automatic Registration Toolbox (ART) (www.nitrc.org/projects/art) resulting in a binary intra-cranial space mask which we denote by *U*^(*m*)^: ℝ^3^ × {1} → {0,1}, where:

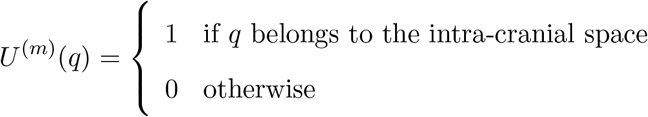

Next, we obtained PIL transformation matrices 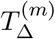 for *V*^(*m*)^ using the procedure described in Section 2.1. Finally, an intra-cranial space probability map was obtained by applying the transformations 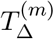 to the binary masks *U*^(*m*)^ and averaging over all *M* cases as follows:

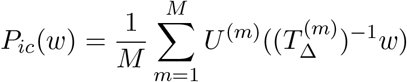

The intra-cranial space probability map *P*_*ic*_: ℝ^3^ × {1} → [0,1] was saved as an auxiliary volume. During the registration process, after transformation of a given volume to PIL space, ATRA recalls *P*_*ic*_ and limits landmark identifications to regions where the intra-cranial probability map values are greater than a prescribed level as shown in Fig. 4.

**Figure 4:**
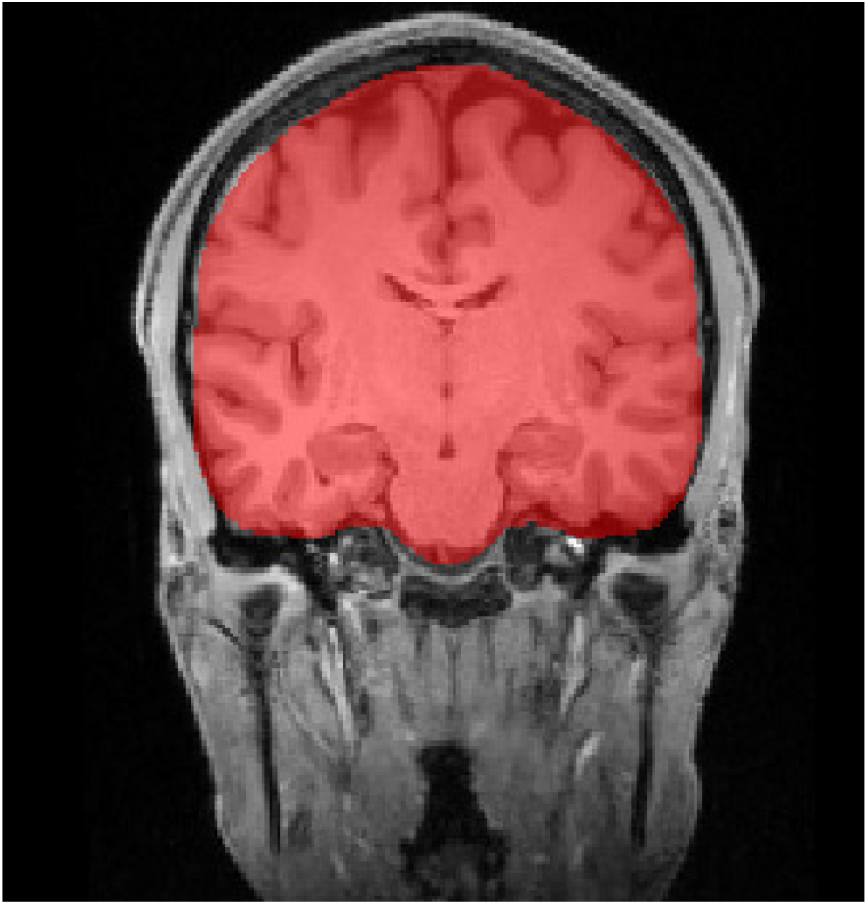
Regions where the intra-cranial probability map *P*_*ic*_(*w*) > 0.5 are shown in red superimposed on a coronal slice of a volume in PIL space. This method is used to to remove almost all non-brain regions from the analyses.

### 2.3 Leave-one-out consistent (LOOC) landmarks

Consider a *supervised* landmark detection algorithm 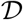, which given a set of *M* example corresponding landmarks *Q* = {*q*^(1)^, *q*^(2)^,…, *q*^(*M*)^} identified on *M training* volumes 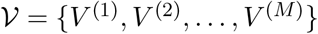, estimates the corresponding landmark *q*^(*t*)^ on a given test volume 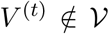. The nature of the landmark detection algorithm is not important for the discussion in this section. The particular algorithm used in ATRA is described in Section 2.5. Other examples are described in Ardekani and Bachman (2009) and Ghayoor et al. (2017). If we denote the estimated landmark on *V*^(*t*)^ as 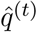, the landmark detection algorithm can be thought of as a function of *V*^(*t*)^ as well as training data *Q* and 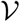 such that 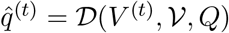.

Next, consider the case where we remove the *m*^th^ training volume *V*^(*m*)^ from the training set 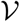 and, accordingly, the *m*^th^ landmark from *Q*. Let 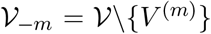 and *Q*_–*m*_ = *Q*\{*q*^(*m*)^} denote the resulting reduced sets, where the notation *X\Y* indicates the relative complement of set *Y* in set *X*. For example, 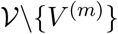 means the set of all elements of 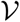 except *V*^(*m*)^, which for short we denote by 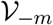.

Now we let our landmark detection algorithm 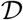 estimate the left-out landmark point *q*^(*m*)^ on the left-out volume *V*^(*m*)^ based in the remaining *M* – 1 training data pairs 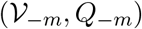. Using the notation above, the estimated landmark is given by 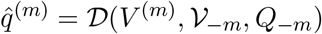.

**Definition.** *A set of corresponding landmarks Q* = {*q*^(1)^, *q*^(2)^,…, *q*^(*M*)^} *identified on volumes* 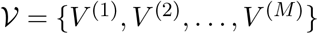 *are leave-one-out consistent (LOOC) with respect to landmark detection algorithm* 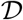 *if and only if* 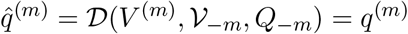 *for all m*.

Thus, to determine whether a set of landmarks are LOOC, we compute the leave-one-out estimates 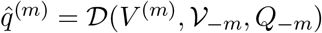 and check to see if 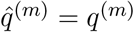 for all *m* ∈ {1,2,…, *M*}. If this is true, then by the above definition, the set of landmarks *Q* is LOOC with respect to algorithm 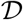.

### 2.4 LOOC landmark identification

Supervised landmark detection algorithms such as the one described in Section 2.5 rely on a set of example landmarks {*q*^(1)^, *q*^(2)^,…, *q*^(*M*)^} identified on a set of *training* volumes {*V*^(1)^, *V*^(2)^,…, *V*^(*M*)^}. Often these landmarks are identified manually by expert raters (Ardekani and Bachman, 2009; Ghayoor et al., 2017). This process, however, is tedious and time-consuming, and suffers from poor inter and intra-rater reliability. Furthermore, we have noticed that manually identified landmarks are almost never LOOC. Also, the manual identification requirement precludes their use in landmark-based automatic registration algorithms such as ATRA. In addition, what humans consider a “good landmark” is not necessarily ideal for machine learning.

In this section, we present a simple but extremely useful algorithm for fully automatic identification of LOOC landmarks that can then be used to train landmark detection algorithms and for automatic image registration. Suppose that a set of training volumes 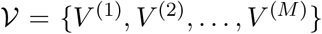 are given after having been transformed to a standard space, for example, to PIL orientation by using the procedure outlined in Section 2.1. Since these volumes are in a standard space, a given point *s* in the standard space *roughly* corresponds to the same anatomical location on all *V*^(*m*)^. The LOOC landmark identification algorithm proceeds as follows:

1. Initially set *q*^(*m*)^ = *s* for all *m* ∈ {1, 2,…, *M*}. That is, at the begining, all landmarks are set to the *same* seed position *s* in the standard space.
2. Repeat the following steps until convergence or until a predefined maximum number of iterations is reached:

2.1. Estimate the leave-one-out predictions 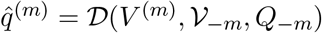. If the landmarks are LOOC, that is, 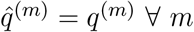, then set a convergence flag and go to step 3.
2.2. If the maximum number of iterations has been reached, then set a non-convergence flag and go to step 3.
2.3. Set 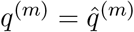 and return step 2.1.
3. End.

If the above algorithm converges within the given maximum number of iterations, then a set of LOOC landmarks has been identified. If convergence is not reached within the prescribed number of iterations, then the algorithm may be repeated with a new seed point *s*. In general, the seed point *s* can be varied systematically and the above algorithm repeated to find multiple sets of LOOC landmarks on the same set of training volumes 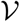. These can be used to train a supervised landmark detection algorithm, such as the one described in Section 2.5, to detect the corresponding landmark on a new test volume *V*^(*t*)^. This is the method we used for training ATRA to detect the 8 mid-sagittal landmarks used for finding *T_lm_* in Section 2.1 (Fig.3). For this purpose, the *V*^(*m*)^ were 30 volumes acquired from different individuals. In addition, if many sets of LOOC landmarks are found on longitudinal volumes, that is, when *V*^(*m*)^ are acquired from the same individual over time, then they can be used for symmetric intra-subject registration as described in Section 2.7.

### 2.5 Landmark detection

Assume that we have a set of *M* training volumes 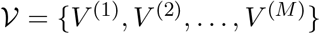 in a standard space and that the location of a given landmark is known on each volume. Let these points be represented by set *Q* = {*q*^(1)^, *q*^(2)^,…, *q*^(*M*)^} where *q*^(*m*)^ denotes the location of the landmark on the *m*^th^ training volume. Points *q*^(*m*)^ can be manually identified on *V*^(*m*)^ by a trained observer, or identified automatically using the method described in Section 2.4. For now, we just assume that *q*^(*m*)^ are known.

This section describes a method for estimating the location of the landmark on a new test volume *V*^(*t*)^ also in the standard space given the training data 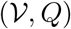. The landmark detection algorithm can be thought of as a function 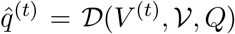, where 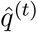 denotes the estimated location of the landmark on the test volume *V*^(*t*)^.

Based on the training data *Q*, the expected location of the landmark in the standard space can be estimated by averaging *q*^(*m*)^:

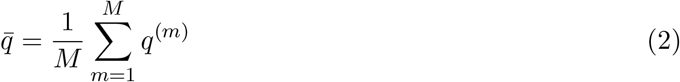

Let 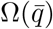 denote a spherical neighborhood centered at point 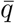. The landmark detection algorithm assumes that 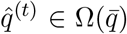, that is, the location of the landmark on the test volume is within some radius of the average location 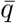.

Furthermore, the landmark detection algorithm relies on a *feature vector* that can be obtained for any point *q* on a volume *V*. The feature vector can be considered a vector-valued function *f* (*V, q*) ∈ ℝ*^l^* that defines an *l*-dimensional vector extracted from volume *V* at point *q*. Given these definitions, the landmark detection algorithm can be written as:

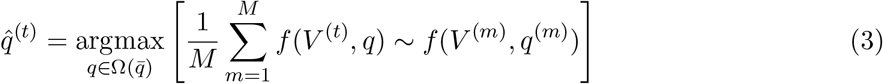

where *f*(*V*^(*t*)^,*q*) ~ *f*(*V*^(*m*)^,*q*^(*m*)^) denotes the degree of *similarity* of the feature vector at point *q* on the test volume *V*^(*t*)^ and the feature vector at point *q*^(*m*)^ on the training volume *V*^(*m*)^. Thus, the landmark detection algorithm (3) calculates the average similarity between *f*(*V*^(*t*)^,*q*) and *f*(*V*^(*m*)^, *q*^(*m*)^) for *m* = 1,2,…, *M* and estimates the landmark on the test volume to be the point in the neighborhood 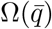 that maximizes the average similarity.

In our current implementation, as feature vector *f*(*V, q*), we simply take the gray levels of *V* in a spherical neighborhood *ω*(*q*) of point *q* and normalize them so that the elements of vector *f*(*V, q*) have zero mean and unit variance. It is possible to include other information in the feature vector function *f*(*V, q*), for example, intensity gradient and/or Laplacian values measured at different resolution levels. However, this line of research has not been pursued in the current work.

As the similarity measure “~” in (3) we simply use the dot product between the feature vectors *f*(*V*^(*t*)^, *q*) and *f*(*V*^(*m*)^, *q*^(*m*)^) which measures the degree of *alignment* between them. The advantage of this choice is that, because the dot product is a linear operation, (3) simplifies to:

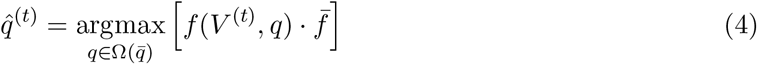

where

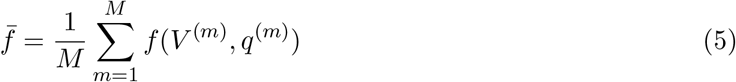

The computational advantage of (4) is that it depends on the *average* feature vector (5) obtained from the training set. The average feature vector 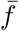 can be computed offline once along with the average landmark location 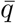. These quantities can then be recalled during landmark detection as auxiliary inputs to the supervised landmark detection algorithm and used in (4) in a computationally efficient manner to detect the landmark location 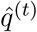 on *V*^(*t*)^.

### 2.6 Generalized Orthogonal Procrustes Analysis

Assume that we have a set of *N* landmarks on each of *M* volumes represented by 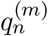 (*n* = 1,2,…, *N*; *m* = 1,2,…, *M*), where 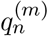 denotes the location of landmark *n* on volume *m* given in terms of its real-world coordinates. The objective of the Generalized Orthogonal Procrustes Analysis (GP) (Devrim, 2003) is to estimate *M* rigid-body transformations 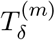, one for each of the *M* volumes, so as to transform them to an average position which is itself unknown. The algorithm is as follows:

1. Compute the *N* centroids 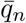 as the average landmark positions across the *M* volumes as follows:

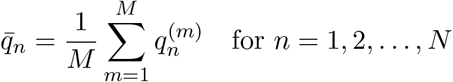
2. Using the algorithm described by Arun et al. (1987) find *M* transformations 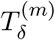 such that

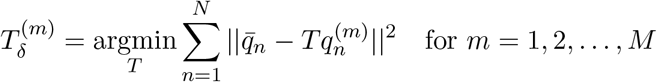

subject to constraints 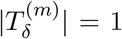. These rigid-body transformations map the *N* landmarks in each volume *m* as closely as possible to the centroids 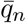.
3. Update 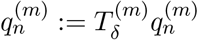 for all *n* and *m*.
4. Repeat steps 1–3 until convergence, when the centroid 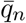, and hence 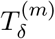, do not change from the previous iteration.

### 2.7 The Automatic Temporal Registration Algorithm (ATRA)

To reiterate, our main objective in this paper is to perform a symmetric registration of a set of volumes 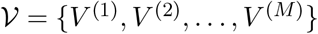 acquired from the same individual over time. For this purpose, we would like to find rigid-body transformations of the form 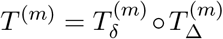 that would register the corresponding volumes to a common space. The transformations 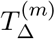 are found individually for each volume using the procedure described in Section 2.1. They transform the volumes into an intermediate PIL space in which the *M* volumes become closely aligned; however, small residual misalignments remain. These small misalignments are corrected by the residual transformations 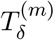 which are found symmetrically using the procedure described in the current section. The ATRA algorithm is as follows:

1. Determine *M* transformations 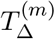 using the method described in Section 2.1.
2. Initialize 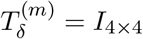, where *I*_4×4_ denotes the 4 × 4 identity matrix.
3. Repeat steps 3.1–3.3 until the number of *anchor points* (*N*\*) as defined in 3.2 reaches a maximum:

3.1 Apply transformations 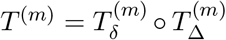 to transform *V*^(*m*)^ to a common space where the set of transformed volumes is denoted by: 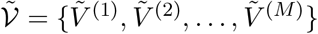.
3.2 Identify *N* sets of LOOC landmarks 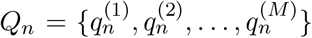 (*n* = 1,2,…, *N*) on 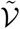 using the LOOC landmark identification algorithm given in Section 2.4. Here, the LOOC landmark identification algorithm is run multiple times using different seed points *s*. The seed points *s* are selected at an equally spaced 3D grid of points in the region *P_ic_*(*s*) > 0.5 (Fig. 4), where *P*_*ic*_ is the intra-cranial space probability map described in Section 2.2. In addition, the seed points s are selected to be symmetric with respect to the MSP. Note that at each iteration, the number *N* may, and usually does, vary depending on the number of seeds that result in convergence to a set of LOOC landmarks. Usually this number increases with increasing number of iterations as the volumes 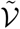 become more closely registered. Consider a set of LOOC landmarks 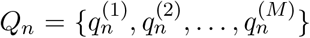 obtained in the current step of the algorithm. If volumes 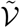 are precisely aligned, then in addition to being LOOC, the landmark locations 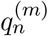 in the common space will likely have the property of being independent of *m*, that is, 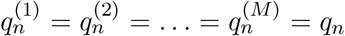. In other words, points 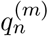 on all *M* volumes 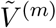 would be represented by the same point *q*_*n*_ in the common space. We refer to these as *anchor points*. Let *N*\* ≤ *N* be equal to the number of identified LOOC landmark sets with this additional property.
3.3 Apply the GP algorithm described in Section 2.6 to the LOOC landmark sets 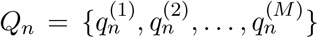 (*n* = 1,2,…, *N*) identified in step 3.2 to obtain a set of updated 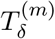.
4. Save transformations 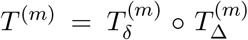 obtained at the iteration where the maximum number of anchor points *N*\* was attained and end.

### 2.8 Practical implementation details

In this section we present the parameters used in our implementation of ATRA which is freely available at www.nitrc.org/projects/art.

#### 2.8.1 Alignment to standard orientation

For PIL transformation (Section 2.1), in the third step of the algorithm in Fig. 2, we find transformation *T_lm_* based on the detection of 8 mid-sagittal landmarks near the MSP. The search centers 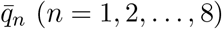 and mean feature vectors

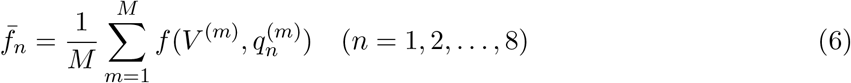

were computed offline and stored in an auxiliary file. This file is uploaded by ATRA during the registration process and used for estimating 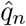 on the volumes being registered. The eight landmarks were found using the LOOC landmark identification algorithm described in Section 2.4 by using *M* = 30 training volumes 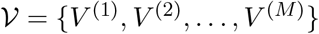 acquired from normal individuals on a 3 Tesla scanner. These volumes were transformed to a standard PIL space using transformations *T_acpc_* ○ *T*_*msp*_. Note that the standard transformations did not include *T*_*lm*_. To identify the 8 sets of LOOC landmarks, multiple seed points *s* were suggested (manually specified) to the LOOC landmark identification algorithm. Then in those runs where the algorithm converged, we visually inspected the landmarks 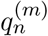 on all 30 volumes to ensure that they point to homologous anatomical points on all brains.

#### 2.8.2 Intra-cranial space probability map

To construct *P*_*ic*_, the intra-cranial space probability map described in Section 2.2, we applied the *brainwash* module of ART to *M* = 152 *T*_1_-weighted volumes obtained from adults of all ages using both 1.5 T and 3T scanners. Minor manual corrections were performed on some of the images before transformation of the binary masks to the PIL space and averaging to obtain *P*_*ic*_. The probability map is distributed with ATRA as an auxiliary volume. In this application, it is used to limit the seed points *s* used by the LOOC landmark detection algorithm to the intra-cranial space. In addition, in the landmark detection algorithm described in Section 2.5, *P*_*ic*_ is used to limit the search region 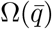 to the intra-cranial space.

#### 2.8.3 LOOC landmark identification

We have defined a set of landmarks to be LOOC when their leave-one-out estimates 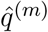 *equal* the left-out values *q*^(*m*)^. In our implementation of the LOOC landmark identification algorithm (Section 2.4), we reorient all images to PIL space and then discretized at a resolution of 1 × 1 × 1 mm^3^. Therefore, in practice, points 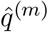 and *q*^(*m*)^ are not in ℝ^3^ × {1}, but belong to a discretized grid and can be described in homogeneous coordinates as [*i j k* 1]^*T*^ belonging to *I* × *J* × *K* × {1} where *I* = {0,1,…, *n_x_* – 1}, *J* = {0,1,…, *n_y_* – 1}, *K* = {0,1,…, *n_z_* – 1}, and *n*_*x*_, *n_y_* and *n_z_* are the number of voxels in the *x, y* and *z* directions, respectively. Therefore, *equality* of 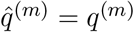 in the context of LOOC landmark identification means that the points have the exact same [*i j k* 1]^*T*^ representation. In our implementation, the default matrix dimensions *n*_*x*_×*n*_*y*_ ×*n*_*z*_ are 255×255×189.

#### 2.8.4 Landmark detection algorithm

The discretization described in Section 2.8.3 also applies to the landmark detection algorithm. In particular, the search represented by Eq. 4 is performed over the discrete domain 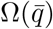 which approximates a spherical search region. In our implementation, the search radius of 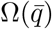 was set to 3 mm (i.e., 3 voxels). The feature vectors *f*(*V, q*) and the average template 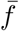 defined in Eq. 5 are also computed in the same discrete domain over the spherical patch *ω*(*q*). In our implementation, the default patch radius of *ω*(*q*) is set to 7mm or (i.e., 7 voxels).

#### 2.8.5 The Automatic Temporal Registration Algorithm (ATRA)

In the final registration algorithm presented in Section 2.7, the LOOC landmark identification algorithm is initialized by multiple seeds *s*. The seeds were selected so that: (1) they are 20 mm apart on a regular 3D grid; (2) they belong to the intra-cranial space with a probability of greater than 0.5, i.e, *P*_*ic*_(*s*) > 0.5); and (3) they are symmetric with respect to the MSP. With these conditions, ATRA starts with a fixed set of 188 seeds points *s*. However, the number of seeds for which landmark identification converges (*N*) varies depending on the problem, as does the number of anchor points (*N*\* ≤ *N*).

### 2.9 MRI data

The MRI data that we used in this study for evaluating ATRA were obtained from two differences sources. Data from 503 subjects were obtained from the ADNI Phase 2 (ADNI-2) database (adni.loni.usc.edu). The ADNI was launched in 2003 as a public-private partnership, led by Principal Investigator Michael W. Weiner, MD. The primary goal of ADNI has been to test whether serial MRI, positron emission tomography (PET), other biological markers, and clinical and neuropsychological assessment can be combined to measure the progression of mild cognitive impairment (MCI) and early Alzheimer’s disease (AD). For up-to-date information, see www.adni-info.org.

Four 3D structural *T*_1_-weighted MRI scans, 2 at baseline and 2 at one-year follow-up, were downloaded for each of 503 ADNI-2 subjects, for a total of 2012 volumes. Images had been acquired using a harmonized pulse sequence on 3 Tesla scanners across 42 different imaging centers. At the time of baseline scans, there were 123 cognitively normal (CN) subjects, 63 subjects with AD dementia, 191 subjects diagnosed with early MCI (EMCI), 112 subjects with late MCI (LMCI), and 14 subjects with subjective memory complaints (SMC). Subjects included 233 females and 270 males. Subjects’ ages ranged from 55 to 92.5years (average: 72.8).

To evaluate ATRA’s accuracy and speed in handling relatively long time-series of 40 longitudinal scans, we also downloaded a unique publicly available dataset from Stanford University (Maclaren et al., 2014) comprised of 120 *T*_1_-weighted volumes from 3 subjects (40 volumes/subject) acquired over 20 MRI acquisition sessions using the ADNI protocol on a GE MR750 3 Tesla scanner. Each subject had been scanned twice within each session, with repositioning between the two scans.

### 2.10 Performance evaluation

We registered the 4 volumes within each of the 503 ADNI subjects and the 40 volumes within each of the 3 Stanford subjects. We recorded the execution time of the algorithm in each case and visually inspected the results for accuracy.

The absence of ground truth registration makes it difficult to quantify the accuracy of registration algorithms on real data. Nevertheless, it is possible to devise a procedure by means of which one can state that the average registration accuracy is at least better than a given prescribed amount.

Since the ground truth registrations are unknown in our data, we devised a novel method for showing that ATRA’s registration accuracy is at least better than 0.5 mm translation or 0.5° rotation. In this method, in each of 20 ADNI subjects selected at random (5 CN, 5 EMCI, 5 LMCI, and 5 AD), after registration of the 4 longitudinal volumes by ATRA, we randomly selected one of the 4 volumes, randomly selected one of the 3 axes, and deliberately added either a 0.5 mm translation along the axis or a 0.5° rotation about the axis to the solution transformation before transforming all 4 volumes to the common space. Then, while being blind to the identity of the deliberately misaligned volume, we attempted to identify it by visual inspection of all 4 volumes side-by-side in the common space. The idea here is that if the average accuracy of the algorithm is on the order of the prescribed motion, then we will only be able to identify the misaligned volume at chance level with 0.25 probability of success. This experiment could then be considered as a Bernoulli process with the number of successes modeled as a binomial distribution B(20, 0.25). If the number of successes achieved in this experiment is significantly larger than the expected value of 4 out of 20, one can infer that the accuracy of registration is better than the prescribed motion.

## 3 Results

We first inspected the 2012 ADNI volumes for accuracy of the PIL transformation method described in Section 2.1. The algorithm failed for 44 volumes (2.2%). Twenty-three of the failed cases were due to the failure of the MSP detection algorithm (step 1, Fig. 2). The remaining 21 failed cases were due to the failure of the AC/PC detection algorithm (step 2, Fig.2). There were no failures in the detection of the 8 mid-sagittal landmarks (step 3, Fig. 2). For these cases we manually specified the AC, PC, and a third point (the vertex of the superior pontine sulcus) (VSPS) on the MSP and ran ATRA again while disabling the automatic MSP/AC/PC detection steps, using the manually specified landmarks instead. More specifically, the first two steps in the PIL transformation algorithm (Fig. 2) that finds matrices *T_msp_* and *T_acpc_* were performed using the manually specified AC/PC/VSPS landmarks.

Following this step, the registrations from all 503 subjects were visually inspected by displaying the 4 aligned volumes for each subject side-by-side. The registrations in all cases were deemed to be accurate by visual inspection. The average processing time in these 503 cases was less than 30 seconds running on a MacBook Pro with a 2.7 GHz Intel Core i7 processor.

In quantitative analysis, we attempted to detect a deliberately misaligned volume post-registration by visual inspection, and were able to correctly identify the misaligned volume in 16 out of 20 cases. Under the null hypothesis of a Bernoulli process with 0.25 success probability, the probability of 16 successes is *p* < 10^−6^. An example is shown in Figure 5 in which image (c) was correctly judged to be the misaligned volume. In this case, the misaligned volume could be discerned by examining, for example, the fastigium of the 4th ventricle.

**Figure 5:**
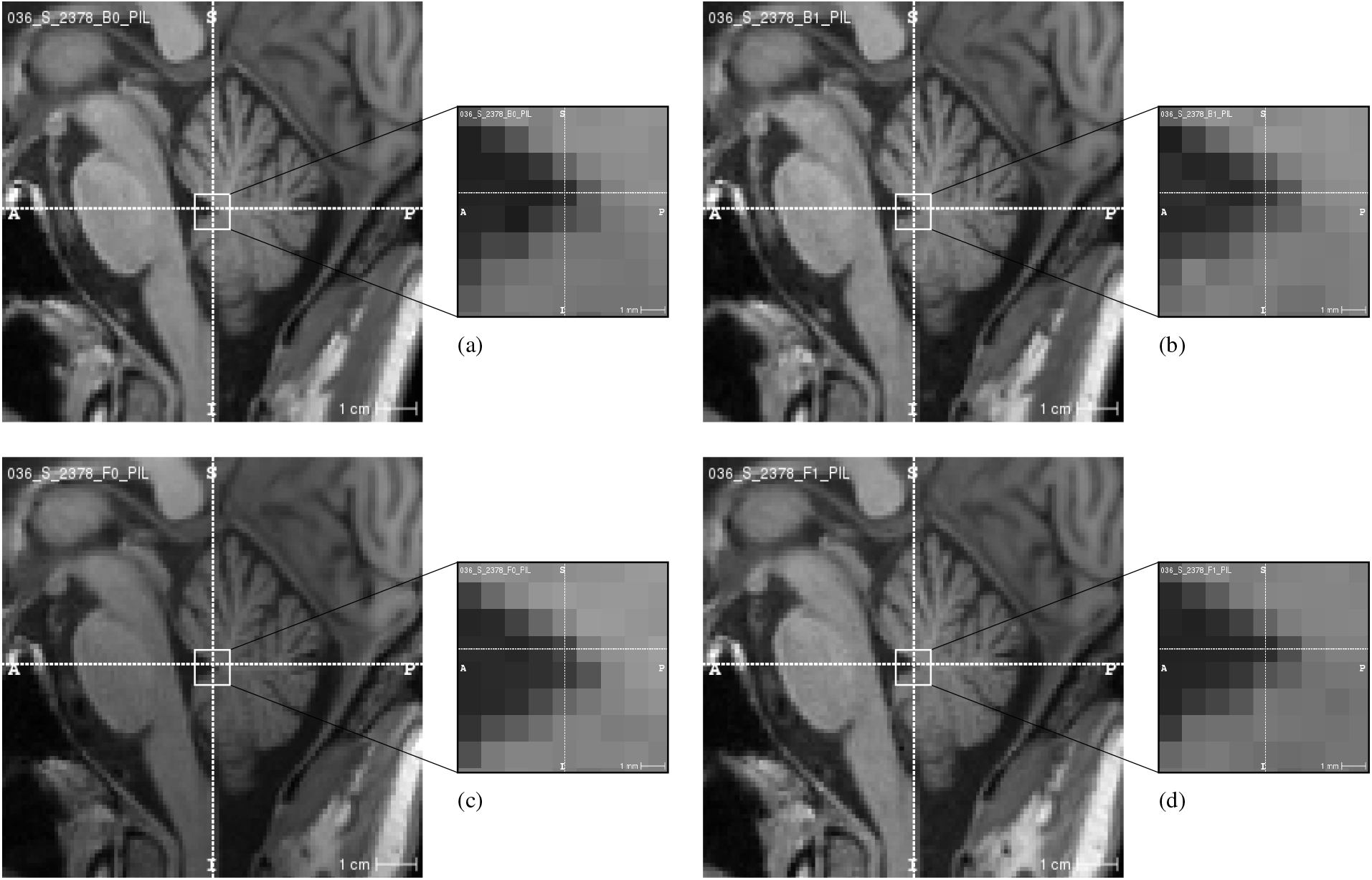
Sagittal sections from 4 longitudinal volumes from an ADNI subject after registration with ATRA. After registration, one of the volumes (c) was deliberately misaligned by adding a 0.5 mm translation to the registration matrix before realignment into a common space. We were then able to distinguish the deliberately misaligned volume by visual inspection, while being blind to its identity. In this example, the misaligned volume could be clearly discerned by examining, for example, the fastigium of the 4th ventricle.

The 40 volumes in each of the 3 Stanford subjects were also registered. There were no MSP or AC/PC detection failures. The post registration aligned volumes were visually inspected and the registrations where deemed to be accurate in all 3 subjects. Figure 6 shows the post registration average of the 40 volumes (right column) in one subject along with a randomly selected post registration volume (volume 20) in the same subject (left column). The average processing time for these cases was 5 minutes and 22 seconds.

**Figure 6:**
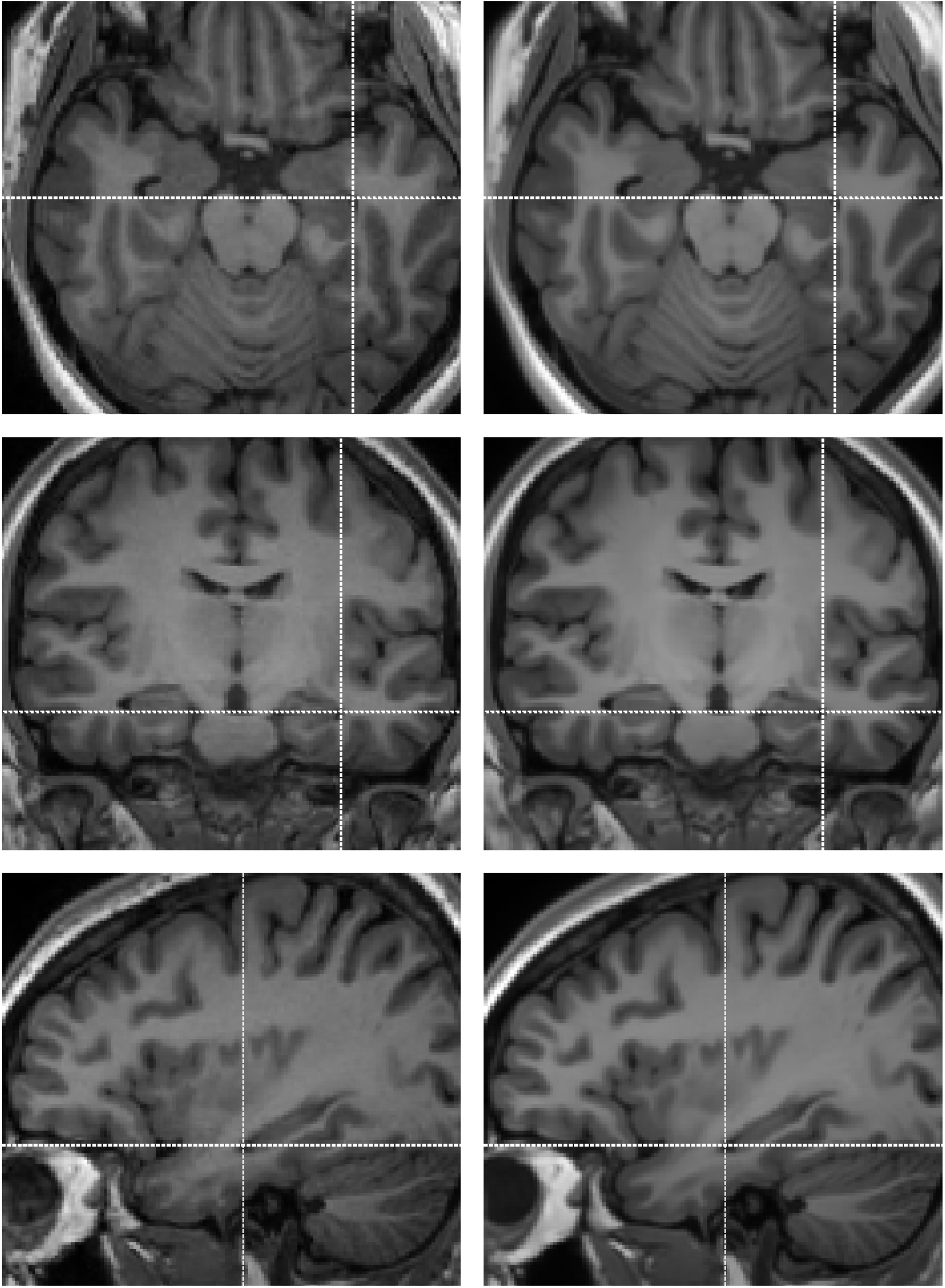
Sections through volume #20 of Stanford subject #3 (left column) and the average of all 40 volumes (right column) after ATRA registration. There is no perceptible loss of resolution in the average images, while the noise level has clearly reduced due to averaging of the 40 volumes.

## 4 Discussion

In this technical report, we presented details of an algorithm for fully automatic symmetric registration of serial 3D *T*_1_-weighted structural MRI scans of the brain obtained from the same individual either as independent scans acquired during the same scanning session or longitudinally over an extended period of time. Along the way, we introduced the notion of LOOC landmarks with respect to a supervised landmark detection algorithm. We also presented a simple but extremely useful algorithm for automatically identifying LOOC landmarks on a set of training volumes. ATRA applies this algorithm to identify a large number of LOOC landmarks across the volumes to be registered; and uses the GP algorithm for aligning the volumes to a common space.

In Section 2.1, we presented an algorithm that can be used to fully automatically realign a given volume to a standard orientation independently based on its intrinsic characteristics. This part of the algorithm is itself a novel contribution to the problem of automatically standardizing the orientation of an MRI scan with numerous applications. Multiple approaches to this problem has been reported in the literature (Arndt et al., 1996; Evans et al., 1992; Li et al., 2003). There are clear advantages of our algorithm over these proposed methods. As an example, Arndt et al. (1996) showed that by locating multiple landmarks on the MSP and using them to estimate a rotation of the MSP into a standard orientation, it is possible to achieve better results than two-point registrations based on AC/PC alone. The third step of our PIL transformation algorithm (Fig. 2) uses a similar idea albeit with several additional innovations. Arndt et al. (1996) relied on manually locating landmarks on the MSP which is time-consuming, requires neuroanatomical knowledge, and can be adversely affected by poor inter-rater reliability. Our method overcomes these issues by detecting the additional MSP landmarks fully automatically. Another innovation of our approach is that we detected the MSP automatically which almost always involves image rotation specified by *T_msp_* in step 1 of the algorithm outlined in Section 2.1, whereas Arndt et al. (1996) select the “middlemost sagittal slice.” Finally, Arndt et al. (1996) use a suboptimal and heuristic method for finding the required translations and rotation of the MSP landmarks to target locations, while we used the procedure described in (Arun et al., 1987) to directly find the *T*_*lm*_ which minimizes the optimality criterion in Eq. (1).

We also formalized an approach for assessing the accuracy of registration whereby one of several volumes post registration is intentionally misaligned by a small prescribed motion. Then, while being blind to which of the volumes was misaligned, we attempt to identify it by visual inspection. If the misaligned volume is correctly identified at a significantly greater rate than what can be expected by chance, then we can conclude that the registration accuracy is better than the prescribed motion. Using this procedure on 20 ADNI cases, we were able to identify 16 cases correctly (*p* < 10^−6^) for a prescribed motion of either a 0.5 mm translation along, or a 0.5° rotation about a randomly selected axis. Thus, we were able to conclude that ATRA achieves sub-millimeter accuracy. It is important to note that the data used for this assessment were from the ADNI database and included subjects with AD (n=5) and MCI (n=10) as well as CN (n=5). We have recently shown (Ardekani et al., 2017) that the brain structure in all these groups undergoes significant atrophy even in the one-year follow-up period of the longitudinal volumes utilized in this work. In many cases, the atrophy results in enlargement of ventricles and expansion of sulci to a degree that is greater than 0.5 mm (half voxel). Nevertheless, we were able to identify such levels of misalignment in these subjects. We attribute this to the presence of *anchor points* in the brain whose relative positions remain stationary over time even though other brain structures may be undergoing changes. ATRA registers longitudinal scans by identifying and aligning a large number of these points.

Our software implementation which has been made freely available relies on setting a number of parameters. These include a 3 mm radius for the search region 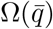 about the expected landmark location 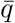, and a 7 mm radius for the spherical patch *ω*(*q*) used to construct the feature vector *f*(*V, q*) at a point *q* on a volume *V*. In addition, in identifying multiple LOOC landmark sets to be used for finding the residual registration matrices *T*_*δ*_ using GP, we used 188 seeds located on a 3D grid of points 20 mm apart in each direction. In the current version of ATRA, none of these parameters was chosen in a systematic way. Changing these parameters from their default value could positively or negatively impact the performance of the algorithm both in terms of accuracy and speed. A systematic study of the impact of these parameters on the algorithm remains a topic of future study.

Decreasing the distance between seeds points *s* from the default value of 20 mm will dramatically increase the number of seed points from 188. Although this is likely to positively affect the accuracy and robustness of the algorithm, it will significantly increase the processing time. On the other hand, the LOOC landmark identification algorithm (Section 2.4), which is the most time-consuming part of ATRA, can be run in parallel for multiple seed points *s*, for example, using multi-core computation. That is, multiple processing units can run in parallel each seeking a set of LOOC landmarks starting from different seed points. Increasing the search area radius from the default value of 3 mm will also increase the processing time but may lead to improvements. Again, to compensate for the additional computation time that will be required to increase the search radius, the search may be accelerated, for example, by GPU computation. Predicting the effect of changing the spherical patch *ω*(*q*) radius is less intuitive. Smaller radii will increase the computation efficiency of the algorithm but may result in deteriorating performance.

We found that the AC/PC or MSP detection algorithm failed in about 2.2% of cases. In all these cases, when we manually supplied to the program three landmarks: the AC, PC, and VSPS, the program proceeded correctly with all subsequent processing steps and resulted in accurate registrations. We used the MSP detection algorithm described in Ardekani et al. (1997) and the AC/PC detection algorithm described in Ardekani and Bachman (2009) because of our familiarity and experience with these methods. Future research may further reduce the already quite small failure rate of these algoirhtms. In addition, other existing algorithms for MSP (Hu and Nowinski, 2003; Prima et al., 2002; Jayasuriya et al., 2013; Volkau et al., 2006) and AC/PC (Bhanu Prakash et al., 2006; Verard et al., 1997; Liu and Dawant, 2015) detection should be evaluated and considered as alternatives if they show reduced failure rates with an acceptable execution time.

A limitation of our current implementation of ATRA is that it was designed for registering *T*_1_-weighted volumes. However, the approach can be extended to other modalities such as *T*_2_-weighted or FLAIR if the training-based methods for MSP, AC/PC and mid-sagittal landmark detection are extended to these modalities. In addition, the ideas may be extended to non-human MRI volumes. Indeed, the automatic LOOC landmark identification algorithm may be applicable to non-medical image processing such as locating landmark points on face images for face recognition.

## Acknowledgement

The author would like to thank professors Alvin H. Bachman and Howard Kushner for helpful discussions and proofreading the manuscript; Mr. Ali Asaei for his help in identifying the 8 mid-sagittal landmarks; Mr. Neema Izadi and Dr. Muhammed Asim Mubeen for their help with downloading and organizing the ADNI data; and Dr. Paul Langman for proofreading the manuscript. Data collection and sharing for this project was funded by the Alzheimer’s Disease Neuroimaging Initiative (ADNI) (National Institutes of Health Grant U01 AG024904) and DOD ADNI (Department of Defense award number W81XWH-12-2-0012). ADNI is funded by the National Institute on Aging, the National Institute of Biomedical Imaging and Bioengineering, and through generous contributions from the following: AbbVie, Alzheimer’s Association; Alzheimer’s Drug Discovery Foundation; Araclon Biotech; BioClinica, Inc.; Biogen; Bristol-Myers Squibb Company; CereSpir, Inc.; Cogstate; Eisai Inc.; Elan Pharmaceuticals, Inc.; Eli Lilly and Company; EuroImmun; F. Hoffmann-La Roche Ltd and its affiliated company Genentech, Inc.; Fujirebio; GE Healthcare; IXICO Ltd.; Janssen Alzheimer Immunotherapy Research & Development, LLC.; Johnson & Johnson Pharmaceutical Research & Development LLC.; Lumosity; Lundbeck; Merck & Co., Inc.; Meso Scale Diagnostics, LLC.; NeuroRx Research; Neurotrack Technologies; Novartis Pharmaceuticals Corporation; Pfizer Inc.; Piramal Imaging; Servier; Takeda Pharmaceutical Company; and Transition Therapeutics. The Canadian Institutes of Health Research is providing funds to support ADNI clinical sites in Canada. Private sector contributions are facilitated by the Foundation for the National Institutes of Health (www.fnih.org). The grantee organization is the Northern California Institute for Research and Education, and the study is coordinated by the Alzheimer’s Therapeutic Research Institute at the University of Southern California. ADNI data are disseminated by the Laboratory for Neuro Imaging at the University of Southern California.

